# Demographic changes of a tropical understory bird in naturally patchy montane habitats in southern India

**DOI:** 10.1101/2021.09.17.460862

**Authors:** V. V. Robin, James E. Hines, James D. Nichols, Madhusudan Katti, Anindya Sinha

## Abstract

The occurrence, density and survival of a species often depend on various aspects of the habitat that it occupies including patch size and disturbance. The demography of most threatened tropical species largely remain unstudied but could provide valuable information about their biology and insights for their conservation. Our study examined the effect of patch size and disturbance on different demographic parameters of an understory, threatened, endemic bird, the white-bellied shortwing in the tropical biodiversity hotspot of the Western Ghats in India. We sampled eight plots on a sky-island using mist-nets for four years in a ‘Robust design’ mark-recapture framework. Based on model selection using AIC values, the model with survival as a function of disturbance fits the data better than models with abundance or with these parameters modeled as functions of forest patch size. Shortwing density and sex ratio were not different across forest patch sizes or differing disturbance regimes. However, the survival rate of the species significantly decreased with increasing disturbance but was not related to forest patch size. Our study is possibly the first to provide critical baseline information on the demography of a tropical understory species from this region.

## INTRODUCTION

One of the most important conservation challenges of our times is the anthropogenic fragmentation of contiguous habitat to form disjunct patches. In this scenario, it has often been argued that preserving large tracts of habitats is essential for the long-term survival of species (Diamond et al. 1987, Beier et al. 2002) though smaller patches could also contribute in this regard (e.g., Githiru and Lens 2006a). In contrast to habitat loss, disturbance unobtrusively affects the demography of different species, especially understory insectivorous birds that are extremely sensitive to disturbances in forest structure (Thiollay 1999, Sekercioglu et al. 2002).

The quality of the habitat can affect critical demographic parameters of a population in various ways. Better-quality areas, for example, can support higher densities of a species due to the greater availability of resources leading to smaller territories (e.g., Githiru and Lens 2006b). Species that are sensitive to fragmentation can also occur less commonly in sub-optimal habitats such as small degraded fragments (e.g., Zanette et al. 2000). Conservation managers, in fact, often use density as a surrogate for the health of a population (e.g., Irwin et al. 2005). However, the rate of turnover in suboptimal habitats can be higher than that in better-quality patches (e.g., Zanette 2001).

Apart from survival rates and population densities, one of the important demographic factors is adult sex ratio (ASR), an indicator of a population’s trajectory, behavioral ecology and its conservation status (reviewed in Donald 2007). ASR is often known to correlate with population trends or habitat quality (Zanette 2001) and has already been used as an indicator of population status in the management of some mammals (Donald 2007). Most research providing information on survival rates and population densities have, however, come from temperate regions.

Tropics, in general, are thought to be inhabited by species that have a “slow” life-history strategy, where high survival rates in optimal patches generate only few openings for new recruits, leading to low individual turnover rates (e.g., Morton and Stutchbury 2000). While there have been a few studies estimating survival rates, particularly using mark-recapture methods, from the tropics (e.g., Karr et al. 1990, Brawn et al. 1995, Brawn et al. 1999, Githiru and Lens 2006b), there have been no studies from the Asian tropics, though they host several global biodiversity hotspots and their endemic species.

The Shola forests in the high-elevation sky islands of the Western Ghats consist of naturally occurring patches of forests that range from 0.1 ha to thousands of hectares, with this patchiness originating from a combination of factors including the geography of the landscape, wind direction and rainfall (Caner et al. 2007). Such patchiness is known to affect various aspects of species demography in both naturally and artificially fragmented landscapes (e.g., Dooley Jr and Bowers 1998). We investigated the effect of such patchiness and habitat disturbance or degradation on the threatened (BirdLife International 2001), endemic, understory insectivorous bird, the white-bellied shortwing *Brachypteryx major* (Robin et al. 2010), also known as white-bellied blue robin. Specifically, we examined the effect of patch size and disturbance on (a) the population density, (b) survival and (c) sex ratios of shortwing individuals in our study patches.

## METHODS

### Study area and design

This study was conducted in the Grasshills National Park, located within one of the sky-islands of the Western Ghats, from 2003 to 2007. This region, lying on a high-elevation plateau, ranging from 1400 m to 2400 m above mean sea level (a.s.l), has an undulating topography in a matrix of natural patches of forests and grasslands (described in Shanker and Sukumar 1998).

We established eight study plots in the Park, at an elevation of 1450 m to 1600 m a.s.l. In one large patch of forest (> 2500 ha), we laid four plots (Plots A to D) of 2.5 ha each, along a disturbance gradient. Disturbance in this landscape resulted entirely from anthropogenic activities, such as the collection of fuelwood. This was subjectively quantified (see below) in different areas before the selection of plots. We selected Plots A to D with a gradient of disturbance decreasing from Plot A to Plot C and with Plot D being an undisturbed plot. We also chose four small, independent, undisturbed patches (Plots E to H; 1.6, 1.1, 0.6 and 0.6 ha in size respectively), where each patch formed a separate plot. Plots E to H were selected such that they would be close to one another and were also the patches closest to the large patch (Plots A to D). This design was adopted to detect any metapopulation-level inter-patch movement, as well as to include patches that varied in size. Note that this selection of study plots, limited by its natural availability, permits some inference about relationships involving disturbance in large continuous habitats, and about relationships involving patch size in relative undisturbed areas, but not inferences about the potential interaction of disturbance and patch size.

### Trapping design

Mist-nets, 12 m × 2 m, at a net density of 10 nets per hectare in each plot, were kept open from the break of dawn (0550 – 0600) for five hours each day, for three consecutive days to capture birds. Each captured bird was tagged with numbered metal bands. The study was designed to be conducted according to Pollock’s (Pollock 1982) Robust Design where primary sampling periods (five) each include multiple secondary sampling periods (three). This design assumes that the population is ‘open’ to immigration, emigration, births and mortality across the four consecutive years (i.e. five primary sampling occasions) in our study while it is ‘closed’ across the three days of secondary sampling occasions within each year. This allowed us to compute the survival rate of individuals in each plot across the primary sampling periods and estimate abundance in the secondary sampling periods.

### Density and survival analysis

Two plot-specific measures, used as covariates of survival rate and abundance, were plot size (a continuous variable, measured in ha) and disturbance category (a categorical variable). The plots were classified into four qualitative disturbance categories, scaling from 0 (undisturbed) to 4 (highly disturbed), based on the number of people encountered, number of cut stems and openness of the canopy.

From the capture data from each plot, we constructed individual capture histories with the ‘1’s indicating captures and ‘0’s indicating no captures on each sampling occasion during each year. Initially, we had planned to use multistate capture-recapture models in order to estimate movement among patches, but the observed numbers of movements was so small (see below) that we instead used single-site models. All juveniles and sub-adults were removed from the analysis. Pollock’s Robust Design originally used closed population models to estimate abundance and the traditional Cormack-Jolly-Seber model (CJS) to estimate survival. The closed population models include estimators of capture probability and abundance that are robust to heterogeneity in detection probability while the CJS model includes survival estimators that are robust to this heterogeneity. Hence, Pollock’s Robust Design provides a suitable framework for estimating both survival and abundance. Kendall et al. (1995) later developed a full-likelihood framework that modeled within- and between-period information simultaneously, permitting the use of reduced parameterizations. We used the program MARK (White and Burnham 1999) to analyze the data using this integrated approach. It should be noted that here ‘survival’ is really “apparent” survival, the complement consisting of mortality as well as permanent emigration, which cannot be separated in this analysis. In the different models, capture and recapture probabilities were modeled as either constant or variable across secondary periods and as variable across primary periods. Individual heterogeneity in capture probability was included using a 2-point mixture (Pledger 2000), which was allowed to vary across the years. Survival, similarly, was modeled as either constant or variable across the four intervals, and/or as differing between study plots. Temporary emigration was modeled as Markovian (γ’, γ’’), random (γ’= γ’’), or absent (γ’= γ’’=0) (Kendall et al. 1997). Additionally, we modeled survival probabilities as functions of patch size and disturbance using the design matrix to specify linear-logistic models. The effects of patch size and disturbance on survival were modeled as interacting with temporal variation or as additive to temporal variation. In all, our candidate set consisted of 39 models. The performance of different models was assessed based on the lowest Akaike Information Criterion (AIC) value. Density was estimated by calculating the effective sampling area by adding a strip width equal to half the average territory diameter to the sides of a plot when similar vegetation continued on that side (Wilson and Anderson 1985).

### Molecular sexing and sex determination

The sexually monomorphic shortwing was sexed using molecular methods with standard primers, P2/P8 (Griffiths et al. 1998) and using a Discriminant Function Analysis (DFA) based on morphometric measurements of wing, tail and beak. Sex was assigned for 91 different individuals of the total 116 captured in this study (described in Robin et al. 2011). Some individuals (25) from across different plots and years remained unsexed and were not used in this analysis as they were without both genetic samples and a complete set of morphological measurements for the DFA to predict their sex. We calculated the sex ratio as the proportion of females in the plot. We combined sex data from all small plots and denoted this combined group as ‘S’ as the small plots E, F, G and H had very few individuals, with some years even characterized by single individuals,. We then examined year- and plot-wise breakdowns of sex ratios and compared them using the Kruskal-Wallis chi-square test in the statistical programming language R (R Development Core Team 2011).

## RESULTS

### Captures and movement

We captured 116 individual adult Shortwings across the four years of sampling. We detected only two instances of movement during the entire study period, one individual from Plot B to A, the most disturbed plots, and another between two small patches (G and H). We were unable to detect any regular patterns of movement across any other plots during this study period.

### Density and survival rate

Models with survival varying as a function of disturbance received the greatest support (AICc weight 0.95985) from the data of all 39 models tested, performing better than models that included plot size (AICc weight 0.22108, Appendix 1). In this model, initial capture probability was constant each year within a season but varied across years, while recapture probability was constant within and across years (Appendix1). Parameter estimates indicated that there was severe trap shyness in the birds, as the estimated recapture probability (0.21 ± 0.03 SE) was much lower than the capture probabilities in different years (0.45 ± 0.04, 0.62 ± 0.05, 1-SE not estimable). Temporary emigration was assumed to be random (Kendall et al. 1997) in the highest ranked models.

The mean population density ± SE across all plots was 2.8 ± 0.37 birds per ha. The abundance estimates varied across years and across plots, and showed that the large patch had a higher abundance of birds than did the small plots. The undisturbed plot had the highest abundance estimate, while all small patches had low abundance estimates. The population density estimates (Figure 1), however, did not show evidence of a difference in density between plots based either on disturbance or size. The temporal variability in the density of the small patches still appeared to be higher than that of the plots in the large patch. Plot C, with very little disturbance, had the lowest estimated density of shortwings. The arithmetic mean survival of the population was 0.58 ± 0.07, which indicated that over half the individuals from the previous year survived to the next year. Survival rate was a function of disturbance in the best model (Figure 2), with disturbed plots having significantly lower survival rates; Plot A, with the highest disturbance level, had the lowest survival rate.

**Figure 1:**
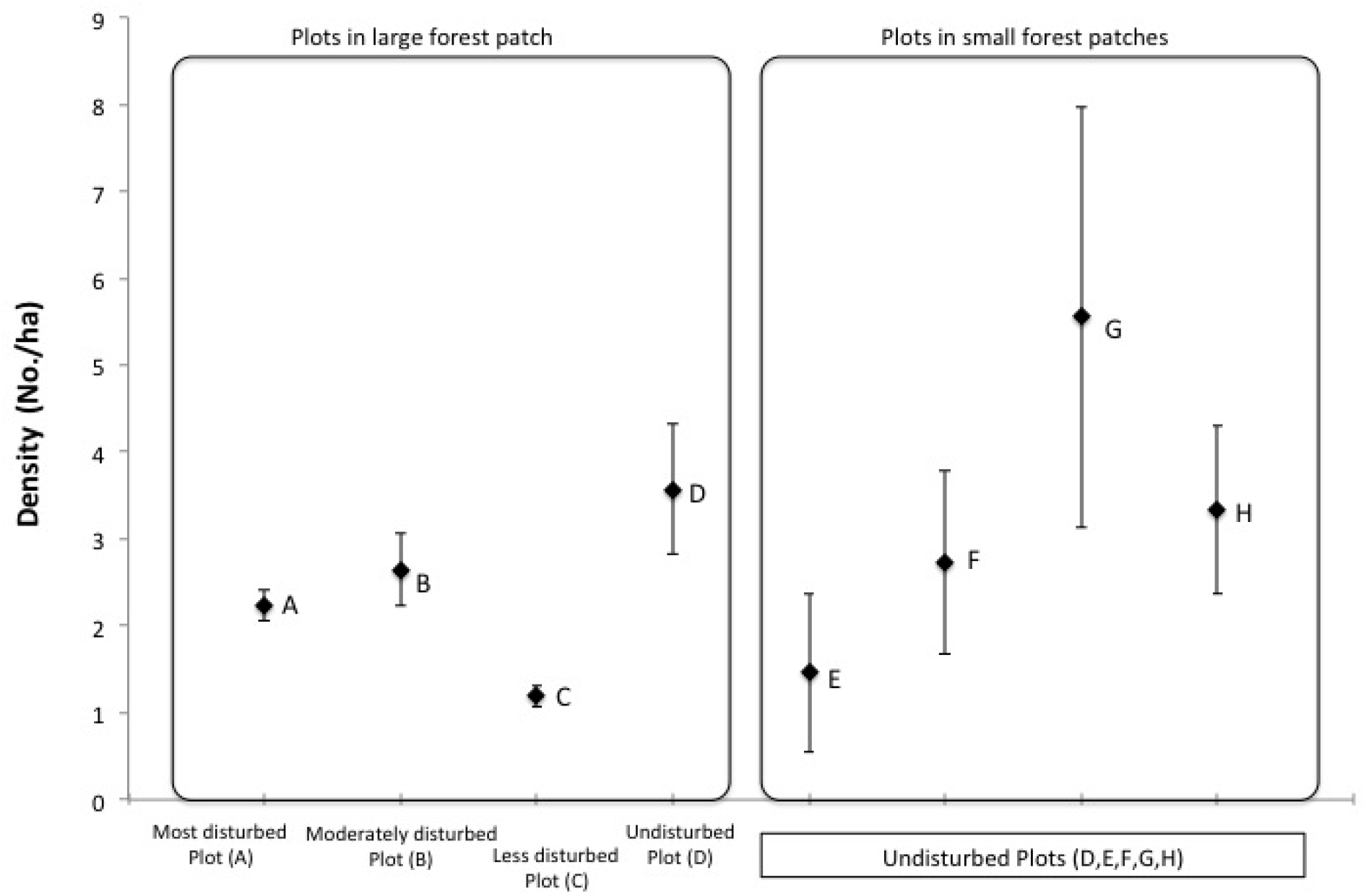
Density of shortwings (number of individuals per ha) in plots across disturbance categories and patch size. Error bars represent SE from yearly variation in density.

**Figure 2:**
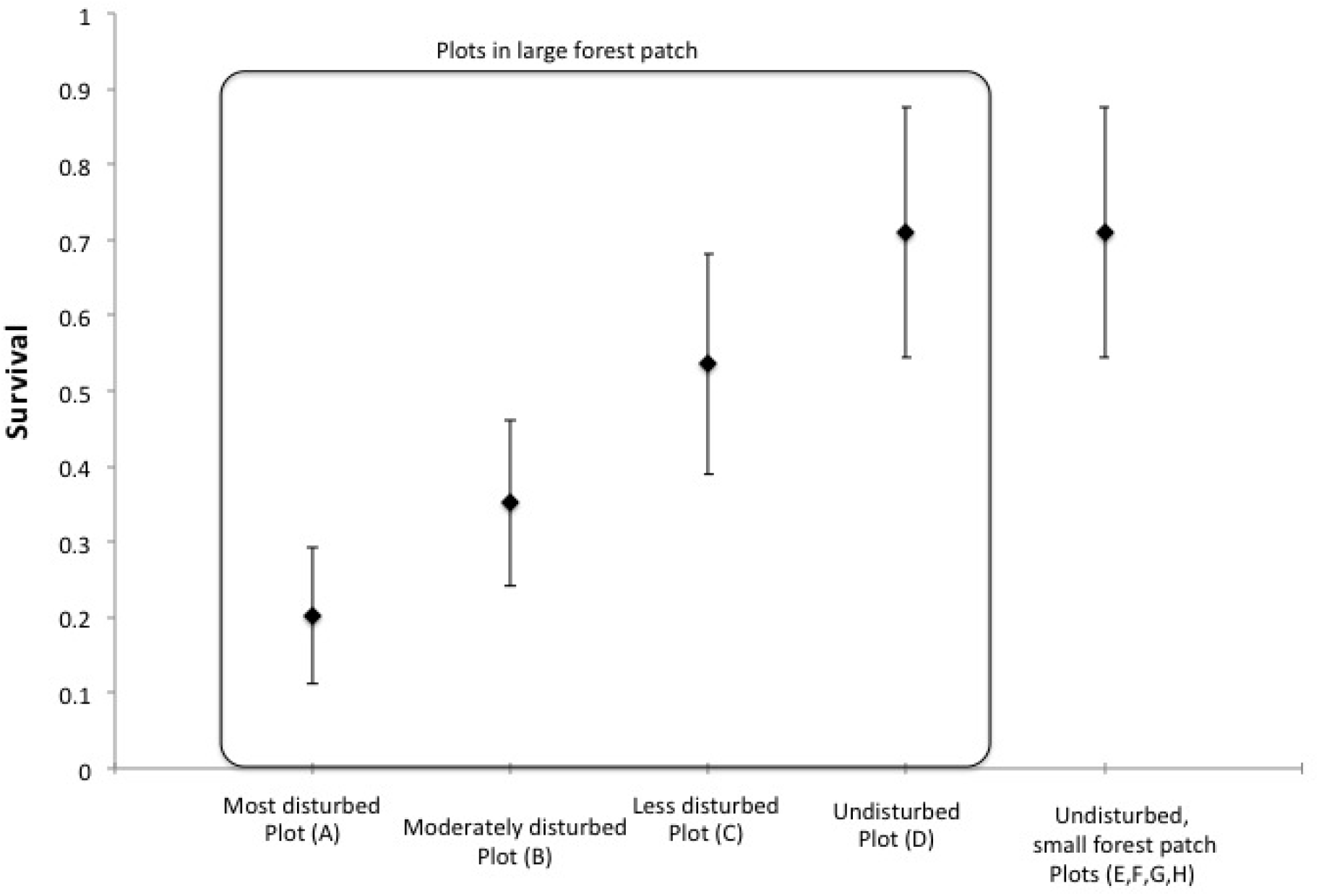
Survival rate of shortwings (expected proportion of individuals that survive and do not permanently emigrate from one sampling year to the next) in plots across disturbance categories and patch size. Note that the plots are aligned along a disturbance gradient, with Plot A being the most disturbed and Plots D to H are undisturbed.

### Adult sex ratio

The overall sex ratio (proportion females) of the entire shortwing population that was sampled across different plots over all years was male-biased (0.44). However, a closer inspection of the data, plot- and year-wise, showed that, barring a few years, the most disturbed plot (A) appeared to be more female-biased than were less disturbed plots (Figure 3). This pattern, however, was not significant over all years (Kruskal-Wallis chi-square = 2.7, df = 2, p = 0.26).

**Figure 3:**
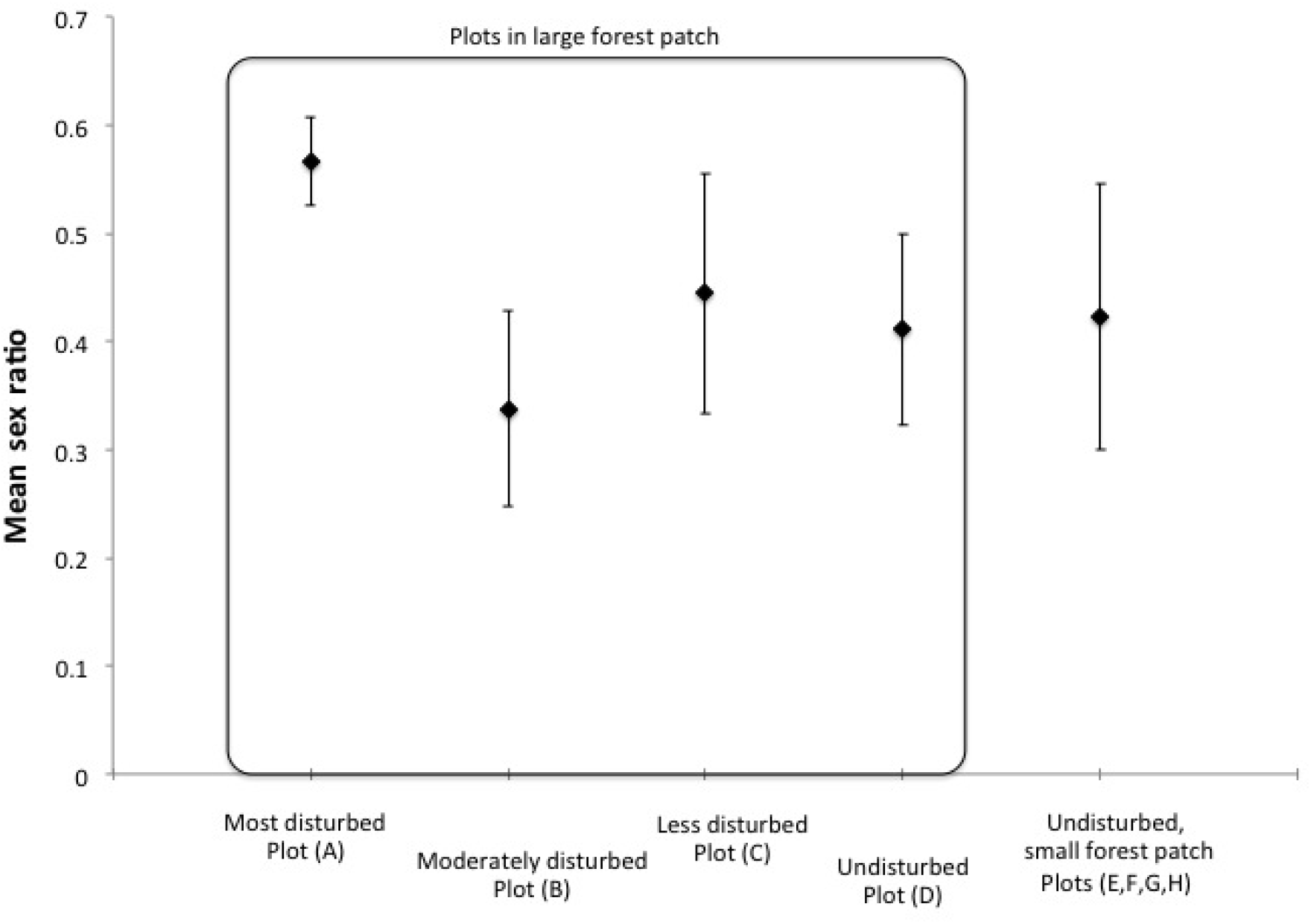
Mean Adult Sex Ratio - Proportion of females in plots. S indicates a sum of all small plots (E to H). Plot A is the most disturbed plot. The error bars indicate standard error across multiple years.

## DISCUSSION

### Density

In general, it is expected that for understory insectivorous birds, fragmented and otherwise sub-optimal habitats harbor a lower density of individuals (e.g., Zanette et al. 2000) compared to the largest and least disturbed fragments (e.g., Githiru and Lens 2006b). The white-bellied shortwing is a species that is highly specific to this habitat and currently considered threatened due to habitat loss. Our results show that there was no consistent difference in shortwing densities across different patch sizes or degrees of disturbance. However, there was a marginally higher estimated density (4.54 individuals/ha) and the highest estimated abundance (14 individuals) of shortwings in the undisturbed plot of the large patch (Plot D). This pattern is similar to that obtained in a study of the white-starred robin, another tropical passerine (Githiru and Lens 2006b). The failure to detect any significant effect of disturbance on shortwing density could be because population density is not sensitive to the changes we are measuring. The lack of baseline data on this species or data on other similar species from this region hinders any further conclusions.

There is very little information on bird densities from the Western Ghats., with Sridhar (e.g., Raman and Sukumar 2002, Sridhar 2005) being the only study with comparable data. This study used point-counts to estimate bird densities in an area that overlapped with one of our study plots (Plot B) during the same study period. The estimated shortwing density of 0.30/ha in one sampling season was approximately ten-fold lower than our estimate for the same year (3.04 ± 0.5) or that averaged over five years (2.6 ± 0.4). This could be because exclusively understory species like the shortwing are inherently difficult to detect in the dense evergreen forest understory, the same reason why the species may have been recorded so few times over the last century (2005).

### Survival rates in small patches

Surprisingly, forest patch size did not affect survival more than did disturbance although the small forest patches were actually very small (< 1 ha). Other studies on similar species such as the white-starred robin (Robin and Sukumar 2002) in Africa also failed to detect an effect of forest patch size on survival although their smallest patch was three times larger (3ha) than that of this study. We believe that the high survival rate in small patches may, once again, be because Shola forest species such as shortwings are adapted to some level of patchiness in the landscape, and individuals consistently persisted in these patches over years. Ancillary data (Robin pers. obs.) indicate that shortwings were using these small patches not just for foraging but also for nesting and breeding, indicating the important role that small patches of natural forest play in the biology and survival of species such as the shortwing.

### Survival in disturbed areas

We found that habitat disturbance was associated with lowered survival rates of shortwings. Our most disturbed plot (A) had, in fact the lowest apparent survival rate of this bird over the entire study period. This could be due to higher mortality rates from predation (Githiru and Lens 2006a), higher levels of permanent emigration, or because these areas are used by a larger number of transients (Gibbs 1991) as has been demonstrated in other tropical regions (Perret et al. 2003). Other studies that have similarly demonstrated higher turnover of individuals in such suboptimal habitats (Johnston et al. 1997). As our data was not sufficient to include transience in our model, we are at present working towards assessing between-patch movement with genetic tools.

### Adult sex ratios

Adult sex ratios (ASR) in wild birds remain very poorly described, though skewed sex ratios are common in the wild (eg. Zanette 2001). Most studies have found male-biased sex ratios in birds while female-skewed ASR appear to be the most common pattern in mammals. We found female-biased ASR in some of our study plots, particularly the most disturbed plot, a rare pattern in birds, having been recorded in only 19 studies on 14 species (9.5% of all studies) so far (reviewed in Donald 2007). Given the generally balanced offspring sex ratio, a skewed ASR can only be explained by higher mortality or dispersal of one of the sexes. Our limited data, with few individuals in each plot, do not permit us to model the effects of transience and sex-specific survival though we plan to do this in the future with additional sampling.

This study has provided evidence that a tropical understory insectivore is adversely affected by habitat disturbance more than by habitat patchiness. Population density did not reflect disturbance effects but survival rate and adult sex ratios were relatively more sensitive to disturbance. Although abundance and population density continue to be used as variables that inform many conservation measures, it appears from our study that other demographic parameters such as survival and adult sex ratio, though less easy to measure, could be more sensitive to habitat disturbance and quality. Anthropogenic habitat fragmentation is conspicuous and has attracted much attention from conservation biologists for its negative effects on population dynamics. In species that have evolved in patchy habitats, where fragmentation predates human influence, more subtle habitat disturbance may impact the survival of such species much more than would larger-scale fragmentation.

## ACKNOWLEDGEMENT

We would like to thank the Tamil Nadu Forest Department for permits and the Ministry of Environment and Forests, Government of India for funding and permits, National Institute for Advanced Studies for support with logistics. D. Jathanna helped with considerable comments and suggestions on this manuscript. A part of the analysis was done while supported by an award from College of Science and Mathematics, California State University, Fresno and with travel support from Patuxent Wildlife Research Centre. We thank R. Nandini, D. Venugopal, E. Vitte, S. Rao, Loganathan, Ramesh, Senthil and Karuppuswamy and Grasshills field staff for help on field.

## REFERENCES

Beier, P., M. Van Drielen, and B. Kankam. 2002. Avifaunal collapse in West African forest fragments. Conservation Biology 16:1097–1111.

BirdLife International. 2001. Threatened birds of Asia: the BirdLife International Red Data Book. BirdLife International, UK.

Brawn, J., J. Karr, and J. Nichols. 1995. Demography of birds in a neotropical forest: effects of allometry, taxonomy, and ecology. Ecology 76:41–51.

Brawn, J., J. Karr, J. Nichols, and W. Robinson. 1999. Demography of forest birds in Panama: how do transients affect estimates of survival rates? Pages 297–305 in Proceedings of the International Ornithological Conference

Caner, L., D. Seen, Y. Gunnell, B. Ramesh, and G. Bourgeon. 2007. Spatial heterogeneity of land cover response to climatic change in the Nilgiri highlands (southern India) since the Last Glacial Maximum. The Holocene 17:195–205.

Diamond, J., K. Bishop, and S. Balen. 1987. Bird survival in an isolated Javan woodland: island or mirror? Conservation Biology 1:132–142.

Donald, P. F. 2007. Adult sex ratios in wild bird populations. Ibis 149:671–692.

Dooley Jr, J. and M. Bowers. 1998. Demographic responses to habitat fragmentation: experimental tests at the landscape and patch scale. Ecology 79:969–980.

Gibbs, J. 1991. Avian nest predation in tropical wet forest: an experimental study. Oikos 60:155–161.

Githiru, M. and L. Lens. 2006a. Annual survival and turnover rates of an Afrotropical robin in a fragmented forest. Biodiversity and Conservation 15:3315–3327.

Githiru, M. and L. Lens. 2006b. Demography of an Afrotropical passerine in a highly fragmented landscape. Animal Conservation 9:21–27.

Griffiths, R., M. C. Double, K. Orr, and R. J. G. Dawson. 1998. A DNA test to sex most birds. Molecular Ecology 7:1071–1075.

Irwin, M., S. Johnson, and P. Wright. 2005. The state of lemur conservation in south-eastern Madagascar: population and habitat assessments for diurnal and cathemeral lemurs using surveys, satellite imagery and GIS. Oryx 39:204–218.

Johnston, J., S. White, W. Peach, and R. Gregory. 1997. Survival rates of tropical and temperate passerines: a Trinidadian perspective. The American Naturalist 150:771–789.

Karr, J., J. Nichols, M. Klimkiewicz, and J. Brawn. 1990. Survival rates of birds of tropical and temperate forests: will the dogma survive? The American Naturalist:277–291.

Kendall, W. L., J. D. Nichols, and J. E. Hines. 1997. Estimating temporary emigration using capture-recapture data with Pollock’s robust design. Ecology 78:563–578.

Kendall, W. L., K. H. Pollock, and C. Brownie. 1995. A likelihood-based approach to capture-recapture estimation of demographic parameters under the robust design. Biometrics 51:293–308.

Morton, E. and B. Stutchbury. 2000. Demography and reproductive success in the Dusky Antbird, a sedentary tropical passerine. Journal of Field Ornithology 71:493–500.

Perret, N., R. Pradel, C. Miaud, O. Grolet, and P. Joly. 2003. Transience, dispersal and survival rates in newt patchy populations. Journal of Animal Ecology 72:567–575.

Pledger, S. 2000. Unified maximum likelihood estimates for closed capture‚Äìrecapture models using mixtures. Biometrics 56:434–442.

Pollock, K. 1982. A capture-recapture design robust to unequal probability of capture. The Journal of Wildlife Management 46:752–757.

Raman, T. and R. Sukumar. 2002. Responses of tropical rainforest birds to abandoned plantations, edges and logged forest in the Western Ghats, India. Animal Conservation 5:201–216.

Robin, V., A. Sinha, and U. Ramakrishnan. 2011. Determining the sex of a monomorphic, threatened, endemic passerine in the sky islands of southern India using molecular and morphometric methods. Current Science 101:676–679.

Robin, V. V., A. Sinha, and U. Ramakrishnan. 2010. Ancient geographical gaps and paleo-climate shape the phylogeography of an endemic bird in the sky islands of southern India. PLoS ONE 5:e13321.

Robin, V. V. and R. Sukumar. 2002. Status and habitat preference of White-bellied Shortwing Brachypteryx major in the Western Ghats (Kerala and Tamilnadu), India. Bird Conservation International 12:335–351.

Sekercioglu, C., P. Ehrlich, G. Daily, D. Aygen, D. Goehring, and R. Sandi. 2002. Disappearance of insectivorous birds from tropical forest fragments. Proceedings of the National Academy of Sciences USA 99:263–267.

Shanker, K. and R. Sukumar. 1998. Community structure and demography of small-mammal populations in insular montane forests in southern India. Oecologia 116:243–251.

Sridhar, H. 2005. Patterns in mixed-species flocking of birds in rainforest fragments of southern Western Ghats. M.Sc. Saurashtra University, Rajkot, Dehradun, India.

Team, R. D. C. 2011. A language and environment for statistical computing. R Foundation for Statistical Computing, Vienna, Austria. ISBN 3-900051-07-0, URL http://www.R-project.org/. Vienna, Austria.

Thiollay, J. 1999. Responses of an avian community to rain forest degradation. Biodiversity and Conservation 8:513–534.

White, G. and K. Burnham. 1999. Program MARK: survival estimation from populations of marked animals. Bird Study 46:120–138.

Wilson, K. R. and D. R. Anderson. 1985. Evaluation of two density estimators of small mammal population size. Journal of Mammalogy 66:13–21.

Zanette, L. 2001. Indicators of habitat quality and the reproductive output of a forest songbird in small and large fragments. Journal of Avian Biology 32:38–46.

Zanette, L., P. Doyle, and S. Tremont. 2000. Food shortage in small fragments: evidence from an area-sensitive passerine. Ecology 81:1654–1666.

