## Appendix Table 1 for "Demographic changes of a tropical understory bird in naturally patchy montane habitats in southern India"

Appendix 1:Models used for White-bellied Shortwing data. p-capture probability, c-recapture probability, s-survival, γ" – emigration parameter, γ’ – immigration parameter, N-abundance, dist- disturbance, t-time and “.”-constant.

| **Model** | **Delta AICc** | **AICc Weight** | **No. Par** | **Deviance** |
| --- | --- | --- | --- | --- |
| {p(t, .) ≠ c(.) s (dist) γ"(.) = γ'(.) N(.)} | 0 | 0.52233 | 8 | 20.65 |
| {p(t, .) ≠ c(.) s (dist + plot size) γ"(.) = γ'(.) N(.)} | 2.17 | 0.17671 | 9 | 20.593 |
| {p(t, .) ≠ c(.) s (dist) γ"(.) ≠ γ'(.) N(.)} | 2.2 | 0.17366 | 9 | 20.628 |
| {p(t, .) ≠ c(t, .) s (dist) γ"(.) ≠ γ'(.) N(.)} | 4.35 | 0.0592 | 10 | 20.528 |
| {p(t, .) ≠ c(t, .) s (plot size + dist) γ"(.) ≠ γ '(.) N(.)} | 6.25 | 0.02291 | 11 | 20.146 |
| {p(t, .) ≠ c(.) s (plot size) γ"(.) = γ' (.) N(.)} | 6.98 | 0.01597 | 8 | 27.626 |
| {p(t, .) ≠ c(.) s (.) γ"(.) = γ'(.) N(.)} | 7.23 | 0.01402 | 7 | 30.083 |
| {p(t, .) ≠ c(.) s (diff plots) γ"(.) = γ'(.) N(.)} | 10.74 | 0.00243 | 14 | 17.62 |
| {p(t, .) ≠ c(t, .) s (.) γ"(.) ≠ γ'(.) N(.)} | 10.77 | 0.00239 | 9 | 29.196 |
| {p(t, .) ≠ c(t, .) s(plot size+dist) γ"(t)= γ'(t) N(.)} | 10.79 | 0.00237 | 13 | 20.037 |
| {p(t, .) ≠ c(t, .) s (plot size) γ"(.) ≠ γ'(.) N(.)} | 10.92 | 0.00222 | 10 | 27.092 |
| {p(t, .) ≠ c(t, .) S(dist) γ"(.)= γ'(.) N(.)} | 11.3 | 0.00183 | 13 | 20.551 |
| {p(t, .) ≠ c(t, .) S(plot size+dist) γ"(.)= γ'(.) N(.)} | 13.61 | 0.00058 | 14 | 20.493 |
| {p(t, .) ≠ c(t, .) s (diff plots) γ"(.) ≠ γ'(.) N(.)} | 15.16 | 0.00027 | 16 | 17.223 |
| {p(t, .)≠c(t, .) s (plot size+dist) γ"(plot)=γ'(plot)N(.)} | 15.24 | 0.00026 | 17 | 14.836 |
| {p(t, .) ≠ c(t, .) S(plot size) γ"(.)= γ'(.) N(t)} | 18.28 | 0.00006 | 13 | 27.526 |
| {p(t, .) ≠ c(t, .) S(diff plots) γ"(.)= γ'(.) N(t)} | 22.93 | 0.00001 | 19 | 17.521 |
| {p(t, .) ≠ c(t, .) s (dist) γ"= γ'=0 N(.)} | 25.82 | 0 | 8 | 46.47 |
| {p(plot size) ≠ c(plot size) s(dist) γ"(.) = γ'(.) N(.)} | 31.95 | 0 | 7 | 54.795 |
| {p(plot size)≠c(plot size) s(dist+plot size) γ"(.)=γ'(.) N(.)} | 36.3 | 0 | 9 | 54.727 |
| {p(t, .) ≠ c(t, .) S(plot size+dist) γ’’= γ’=0 N(t)} | 37.2 | 0 | 13 | 46.449 |
| {p(.) ≠ c(.) s (plot size + dist) γ"(.) = γ'(.) N(.)} | 38.09 | 0 | 7 | 60.935 |
| {p(plot size) ≠ c(plot size) s(dist+plot size) γ"(.) ≠ γ'(.) N(.)} | 38.19 | 0 | 10 | 54.362 |
| {p(.) ≠ c(.) s (plot size + dist) γ"(plot) = γ'(plot) N(.)} | 39.55 | 0 | 10 | 55.728 |
| {p( .) ≠ c(t, .) s (dist) γ"(.) ≠ γ'(.) N(.)} | 39.84 | 0 | 8 | 60.488 |
| {p(.) ≠ c(.) s (plot size + dist) γ"(dist+plot) = γ'(dist+plot) N(.)} | 40.5 | 0 | 9 | 58.922 |
| {p(dist) ≠ c(dist) s(dist+plot size) γ"(.) ≠ γ'(.) N(.)} | 42.61 | 0 | 10 | 58.782 |
| {p(plot size + dist) ≠ c(plot size + dist) s(dist+plot size) γ"(.) ≠ γ'(.) N(.)} | 42.77 | 0 | 12 | 54.35 |
| {p(.) s(plot size+dist) C(.) &≠P γ”= γ’=0 N(t)} | 53.94 | 0 | 10 | 70.114 |
| {P(t, t) ≠ c(t, t) S(plot size+dist) γ”= γ’=0 N(t)} | 64.37 | 0 | 33 | 19.862 |
| {p(.) s(dist) C≠P γ”= γ’=0 N(T*Plot)} | 103.89 | 0 | 31 | 65.439 |
| {p(.) ≠ c(.) s (plot size + dist) γ"(.) = γ'(.) N(t*plot)} | 104.43 | 0 | 33 | 59.92 |
| {p(.)≠ c(.) s(plot size+dist) γ”= γ’=0 N(t*Plot)} | 106.88 | 0 | 32 | 65.42 |
| {p(.) ≠ c(.) s (plot size + dist) γ"(.) ≠ γ'(.) N(t*plot)} | 107.51 | 0 | 34 | 59.91 |
| {p(.) ≠ c(.) s(plot size+dist) γ”= γ’=0 N(t*Plot)} | 109.39 | 0 | 32 | 67.938 |
| {p(.) = c(.) s(dist) γ”= γ’=0 N(T*Plot)} | 120.04 | 0 | 30 | 84.556 |
| {p(.) = c(.) s(Plot Size) γ’=0 N(T*Plot)} | 129.15 | 0 | 30 | 93.669 |
| {p(.) = c(.) s(Plot) γ”= γ’=0 N(T*Plot)} | 137.28 | 0 | 36 | 83.353 |
